# Influence of visual perception on movement decisions by an ungulate prey species

**DOI:** 10.1101/2023.03.10.532105

**Authors:** Blaise A. Newman, Jordan R. Dyal, Karl V. Miller, Michael J. Cherry, Gino J. D’Angelo

## Abstract

Visual perception is dynamic and depends on physiological properties of a species’ visual system and physical characteristics of the environment. White-tailed deer (*Odocoileus virginianus*) are most sensitive to short- and mid-wavelength light (e.g., blue and green). Wavelength enrichment varies spatially and temporally across the landscape. We assessed how the visual perception of deer influences their movement decisions. From August-September 2019, we recorded 10-min locations from 15 GPS collared adult male deer in Central Florida. We used Hidden-Markov models to identify periods of movement by deer and subset these data into three time periods based on temporal changes in light environments. We modeled resource selection during movement using path-selection functions and simulated 10 available paths for every used path. We developed five a priori models and used 10-fold cross validation to assess our top model’s performance for each time period. During the day, deer selected to move through woodland shade, avoided forest shade, and neither selected nor avoided small gaps. At twilight, deer avoided wetlands as cloud cover increased but neither selected nor avoided other cover types. Visual cues and signals are likely more conspicuous to deer in short-wavelength-enriched woodland shade during the day, while at twilight in long-wavelength-enriched wetlands during cloud cover, visual cues are likely less conspicuous. The nocturnal light environment did not influence resource selection and likely has little effect on deer movements because it’s relatively homogenous. Our findings suggest visual perception relative to light environments is likely an underappreciated driver of behaviors and decision-making by an ungulate prey species.

**Summary Statement:** We assessed how visual perception of white-tailed deer influences movement decisions. Our findings suggest visual perception relative to light environments represents an underappreciated driver of decision-making by ungulate prey species.

## INTRODUCTION

The perceptual space of an animal influences decision-making processes and behaviors (Aben et al., 2021; Fagan et al., 2017; Jordan and Ryan, 2015). Prey species rely on perceptual information to navigate heterogeneous landscapes of risk and reward to optimize their probability of survival and reproduction. For many prey species, vision is an important part of their perceptual space and indicates the immediate presence of risk or reward (Cronin et al., 2014). The visual information available to an animal is constrained by the physiological properties of its sensory system and the physical characteristics of its environment. Thus, the visual perception of a prey species depends on its location in time and space (Aben et al., 2021; Endler, 1993). Given the potential tradeoffs associated with movement through different environments and at different times, movement decisions of prey species likely are influenced by visual perception.

Endler (1993) identified five major forest light environments defined by forest geometry, weather (i.e., cloud cover), and solar angle. During the day, light in large gap forests and open areas is enriched across a broad range of wavelengths (white); light in small gaps of forest is long wavelength enriched (yellow-orange); light in woodland shade is short wavelength enriched (blue-gray; Figure 1a); and light in forest shade is middle wavelength enriched (yellow-green; Figure 1a; Endler 1993). At twilight, when the solar angle is < 10° above the horizon, ambient light across the landscape is deficient in middle wavelengths (570-630 nm) resulting in a purple-enriched light environment (Figure 1b). Wavelength enrichment in small gap, woodland shade, and forest shade converge to the white spectrum prevalent in large gap forests and open areas under cloudy weather conditions, while twilight light environments become long-wavelength enriched (yellow to red; Figure 1b) with cloud cover before transitioning to purple. Middle wavelengths also dominate the nocturnal light environment with spectral effects of lunar altitude, lunar phase, and canopy openness resulting in small relative changes in wavelength enrichment across the landscape (Veilleux and Cummings, 2012). Variation in light environments affects color perception and contrast sensitivity which can interact with plant and animal color patterns to make them either more or less conspicuous depending on the transmission and reception of light signals (Endler, 1993; Endler and Théry, 1996). The low-ultraviolet transmission of arctic lichens (Tyler et al., 2014) in the snow-covered and short-wavelength enriched tundra landscape makes them readily detectable to ultraviolet sensitive species like caribou (*Rangifer tarandus*; Hogg et al, 2011; Tyler et al, 2014; Fosbury and Jeffery, 2022). For multiple bird species, ambient light environments play an important role in the evolution of plumage coloration and intraspecific communication (Endler and Théry, 1996; Heindl and Winkler, 2003; Hernández-Palma, 2016). Wire-tailed manakins (*Pipra filicauda*), a lekking species with vivid plumage, frequently display in forest shade which reduces their conspicuousness to predators while maximizing their plumage contrast for intraspecific communication (Heindl and Winkler 2003).

**Figure 1.**
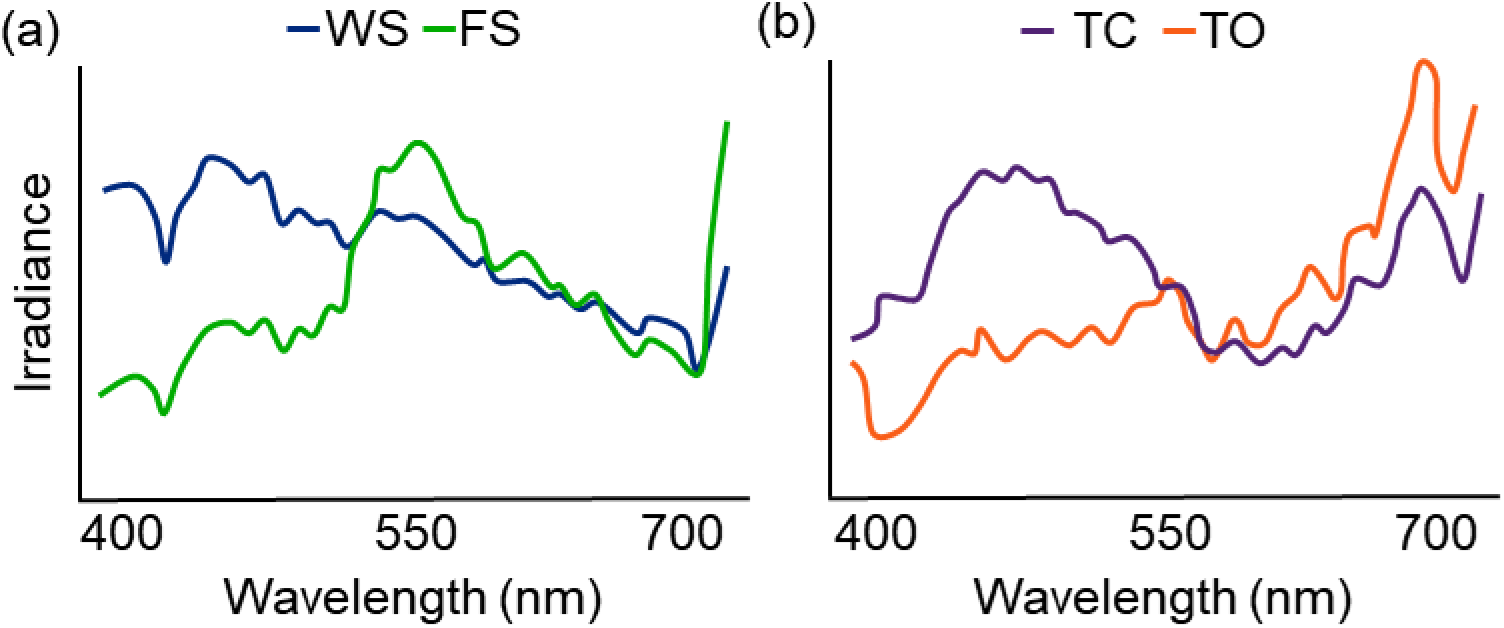
(a) Daytime light environments in woodland shade (WS) and forest shade (FS) based on wavelength enrichment. (b) Twilight light environment during clear conditions (TC) and its changes under cloudy, overcast conditions (TO) based on wavelength enrichment. Mean curves for each light environement were obtained from Endler (1993).

Some light environments have dynamic components that can influence the detection and avoidance of predators by prey (Cuthill et al., 2019). For cryptic species, individual motion negates the benefit of camouflage because movement against a stationary background is highly conspicuous and causes a “pop-out” effect (Rushton et al., 2007). However, dynamic illumination (e.g., dappled forest light) creates non-stationary backgrounds that increase visual complexity and noise within a light environment (Cuthill et al., 2019; Matchette et al., 2019). Matchette et al. (2019) found dappled light masked the movement of prey and increased the time required for predators to visually fixate on prey. Mobile prey might take advantage of these visually noisy and dynamic environments for predator avoidance. Alternatively, prey might avoid dynamic environments because predator movements and cues would also be concealed (Cherry and Barton, 2017). For example, mule deer (*Odocoileus hemionus*) select more open habitats and formed larger aggregations in response to increasing wind speeds likely because of a decreased ability to detect predator movement in visually noisy conditions (Bowyer et al., 2001).

To expand our understanding of how visual perception influences decision-making by an ungulate prey species, we assessed white-tailed deer (*Odocoileus virginianus*) resource selection during movement relative to available light environments. White-tailed deer are a broadly distributed herbivore prey species of the Americas characterized by a reddish-brown to grayish-brown coat with a white underbelly and a distinctive two-tone tail. The countershading of white-tailed deer provides camouflage by reducing shadow in well-lit environments (Caro, 2005; Cuthill et al., 2016), while tail-flagging behaviors might provide a conspicuous signal to maintain group cohesion during flight (D’Angelo et al., 2008) and deter or confuse predatory pursuit (Caro et al., 2020; Loeffler-Henry et al., 2018). Common predators of white-tailed deer include wolves (*Canis* spp.), coyotes (*Canis latrans*), bears (*Ursus* spp.), jaguars (*Panthera onca*), cougars (*Puma concolor*), lynx (*Lynx* spp.), and humans (*Homo sapiens*) (Putnam 1989). Similar to many other mammalian prey species, white-tailed deer possess laterally positioned eyes and a visual streak in their M-cone topography (D’Angelo et al., 2008; Schiviz et al., 2008) that facilitates a panoramic field of view for enhanced predator detection (Banks et al., 2015). Other visual characteristics of white-tailed deer include dichromatic vision with a high perceptual sensitivity for short and middle wavelengths (Cohen et al., 2014; Jacobs et al., 1994) and a reflective tapetum lucidum spectrally tuned for deer photopigments (D’Angelo et al., 2008).

We developed competing hypotheses that white-tailed deer resource selection during movement would be driven by light environment, dynamic illumination, land cover type, or thermal refuge. First, we hypothesized white-tailed deer resource selection during movement would be influenced by forest light environments. During the day, we predicted white-tailed deer would select to move through woodland shade and forest shade while avoiding small gaps in forest because of the relative wavelength enrichment in each habitat and the corresponding spectral sensitivity of white-tailed deer. At twilight, we predicted white-tailed deer selection of forest light environments would be less prevalent (i.e., neither select nor avoid) because of the broad homogenization of the light environment during this time period, while selection or avoidance would suggest factors unrelated to visual perception drive resource selection. At night, we predicted selection of forest light environments during movement would be related to moon illumination. Second, we hypothesized white-tailed deer resource selection during movement would be influenced by dynamic illumination. During the day, we predicted white-tailed deer would alter their selection of light environments as a function of wind speeds to avoid increased visual noise during movement. At night, we predicted no effect of wind speed on light environment selection during movement because of the lower temporal resolution of rods. Support for this hypothesis at night would be indicative of an alternative reason for selection of habitats based on wind speed rather than dynamic illumination. Third, we hypothesized white-tailed deer resource selection during movement would be influenced by land cover types under converging or homogenous light environment conditions. During the day, at twilight, and at night, we predicted movement through land cover types would vary depending on cloud cover. Support of this hypothesis without the interaction with cloud cover would suggest land cover types were more influential than light environments. Finally, we hypothesized the importance of thermal refuge would be more important than light environment. We predicted as temperatures increased deer would select to move through small gaps and forest shade during the day for thermal relief. Results from this study will expand our current understanding of white-tailed deer visual ecology and the influence of light environments and visual perception on movement decisions by an ungulate prey species.

## METHODS

### Study Area

We conducted our study on a private ranch spanning 37,024 ha in Brevard and Osceola counties of Central Florida. Habitat types of the ranch were comprised of improved pasture (62%), freshwater marshes and prairies (17%), hardwood hammocks (8%), flatwoods (5%), cypress domes (4%), and open water (4%; Kawula and Redner 2018). Improved pastures were composed primarily of perennials including bahiagrass (*Paspalum notatum*) and limpograss (*Hemarthria altissima*), as well as planted seasonally for additional winter forage with annuals like ryegrass (*Lolium multiflorum*). The ranch’s wildlife management plan emphasized habitat diversity through edge development and sustainable agricultural operations like rotational grazing.

### Capture and Monitoring

We captured 19 adult (> 1.5 years) male white-tailed deer, hereafter deer, using a helicopter (R-44, Robinson Helicopter Company, Torrance, CA, USA) and net gun during September 2018 (Webb et al., 2008). We minimized pursuit times (<5 min) to reduce capture stress and once restrained, transported deer to work-up stations within 2 km using a sling bag. Processing time was <10 min for every deer during which we estimated the deer’s age using tooth wear, tagged individuals with numbered ear tags (Allflex USA Inc., Dallas, TX, USA), and fitted each deer with a GPS collar (Advanced Telemetry Systems, ATS, Isanti, MN, USA). Once processed, deer were released from the work-up station’s location to limit additional transport stress. Capture and handling protocols were approved by the Institutional Animal Care and Use Committee at the University of Georgia (AUP #: A2018 06-025-R1) and the Florida Fish and Wildlife Conservation Commission (Permit #: SPGS-18-40-A1).

One year later, we programmed the GPS collars of the 15 surviving individuals to record locations every 10 minutes on two separate occasions, the 3-16 of August 2019 and the 24 August-4 September 2019. Aerial surveys were conducted during the 7-9 of August 2019 from 0700-1000. Dyal et al. (2022) found minimal impact of helicopter surveys on deer movement. Thus, we did not censor 10-min locations collected during or after aerial surveys. Locations of animals that experienced mortality during the study were censored following the last fix known to be alive. We recorded an average GPS collar error (± SE) of 4.4 ± 1.1 m in open-canopy areas and 8.7 ± 0.5 m in closed-canopy areas resulting in a cumulative average error of 7.6 ± 0.9 m.

### Statistical Analysis

We obtained 5-min wind and cloud cover data and 1-hr temperature data from an automated station at the Melbourne Regional Airport via the Iowa Environmental Mesonet (available at https://mesonet.agron.iastate.edu/). We converted cloud cover to a continuous factor ranging from clear (0) to overcast (1). We averaged wind speed and cloud cover between 10-min location fixes and used a 1-hr average for temperature. We accessed 30-m resolution land cover and tree canopy data for 2016 from the Multi-Resolution Land Characteristics consortium (available at https://www.mrlc.gov/data). We grouped land cover values into five categories for analysis: cultivated, wetland, developed/barren, forested, and shrub/herbaceous. We reclassified tree canopy values into four forest geometry categories based on the light environments identified by Endler (1993): large gap, < 35% closure; woodland shade, 35-70% closure; small gap, 70-85% closure; and forest shade, > 85% closure. We used the R package *suncalc* (Thieurmel and Elmarhraoui, 2019) to obtain lunar fraction (i.e., the fraction of the moon’s illuminated surface ranging from new moon [0] to full moon [1]) and moon rise and set times for each observation day. Similar to Huck et al. (2017), we used the times of moon rise and set to determine if an observation occurred before moon rise or after moon set. In these cases, the moon fraction was set to 0 regardless of its previous value as the moon would not be visible during these periods. Given temporal differences in deer behavior (Webb et al., 2010) and light environment (Endler, 1993; Veilleux and Cummings, 2012), we subset 10-min locations by time period to analyze separately. Time periods were based on the following criteria: day, period between sunrise and sunset; night, period between sunset and sunrise during which the solar angle is > 10° from the horizon; and twilight, period between sunset and sunrise during which the solar angle is < 10° from the horizon. We used the R package *maptools* (Roger et al., 2019) for sun ephemerides calculations and estimation of time period timings.

We identified movement behavior from the step length and turning angle of each deer using hidden Markov models in the R package *moveHMM* (Michelot et al., 2016). We ran a two-state hidden Markov model which characterized behavior as either encamped (i.e., a state with short steps and wider turning angles) or movement (i.e., a state with longer step lengths and directed movement). To determine the appropriate starting values for step length and turning angle distribution parameters, we compared 25 different parameter sets to ensure models were numerically stable and the final parameter set selected maximized the likelihood function (Michelot et al., 2016). We extracted distance to non-forested wetlands from the Florida Cooperative Land Cover Map (available at https://myfwc.com/research/gis/regional-projects/cooperative-land-cover/) for each location. Based on previous observations of deer behaviors, we expected differences in the transition probabilities between states based on time of day and distance to non-forested wetlands and included these factors as predictors in our final model. We then used the Viterbi algorithm (Zuchini et al., 2016) to identify the most likely behavioral state for each observation based on our model and used only movement state observations for all subsequent analyses. The resulting movement dataset contained 14,658 used paths taken by 15 deer.

We used path-selection functions to model deer resource selection during movement for our three time periods. Path-selection functions provide a fine scale of inference to assess the influence of environmental factors on deer movements by comparing the used path versus available path characteristics (e.g., proportion woodland shade) between sequential movement steps. For every observed path, we simulated 10 available paths drawn at random from a distribution of movement step lengths and turning angles unique to each individual (i.e., 11 points per stratum). We extracted the proportion of our spatial covariates (i.e., land cover type and forest geometry) along a straight line between each used and available path segment using the R package *amt* (Smith et al., 2022). We developed five a priori hypotheses (forest light environment, dynamic illumination, thermal refuge, converging light environment, and a global model) to evaluate deer path selection during movement based on forest geometry, land cover, and weather, as well as moon illumination for night models (Table 1). Because weather variables (i.e., temperature, wind, and cloud cover) and moon illumination were constant within strata, we only included them within our models as interaction terms with spatial covariates. We tested for collinearity among our predictors using a variance inflation factor of < 3. Proportion large gap and proportion cultivated were both highly correlated with multiple other continuous predictors. We removed both proportion large gap and proportion cultivated from their respective models. We fit conditional logistic regressions using the R package *mclogit* (Elff, 2021). We ranked models using Akaike’s Information Criterion (AIC) and selected the most parsimonious model for each time period subset. We identified informative parameter estimates based on whether their upper and lower 85% *CI* overlapped zero (Arnold, 2010). To assess the performance of our top models, we used 10-fold cross-validation (Boyce et al., 2002). We randomly selected 80% of our path data to function as a training set (with 1:10 path strata remaining intact) and allocated the remaining 20% as a test set for the newly trained model. We repeated this procedure nine more times and used the trained model with our test sets to estimate the relative probability of selection of each used or available path. If the proportion of used paths correctly predicted from our pooled test sets was > 0.5, we deemed our model a better fit than would be expected at random. We performed all statistical analyses in R version 4.0.4 (R Core Team, 2021).

**Table 1.** Predictor variables for models predicting path selection for white-tailed deer (*Odocoileus virginianus*) during day, night, and twilight time periods in Central Florida, August – September 2019.

| Time period | Model | Predictors |
| --- | --- | --- |
| Day and Twilight | Forest Light Environment | Forest geometry |
|  | Dynamic Illumination | Forest geometry*wind |
|  | Thermal Refuge | Forest geometry*temperature |
|  | Converging Light Environment | Land cover*cloud cover |
|  | Global | All of the above |
| Night | Forest Light Environment | Forest geometry*moon |
|  | Dynamic Illumination | Forest geometry*wind*moon |
|  | Thermal Refuge | Forest geometry*temperature |
|  | Converging Light Environment | Land cover*cloud cover*moon |
|  | Global | All of the above |

## RESULTS

During movement behavior, white-tailed deer traversed an average of 72 ± 78 m (± SD) in 10 min. The longest observed movement in 10 min spanned 1.4 km and occurred during the day. The forest light environment model was the top model for the daytime period and contained predictors based on forest geometry (Table 2; Figure 2). During the day, deer selected to move through woodland shade (85% CI 0.00, 0.06; SE = 0.02), avoided movement through forest shade (85% CI –0.12, –0.06; SE = 0.02), and neither selected nor avoided movement through small gap (85% CI –0.03, 0.02; SE = 0.02). The converging light environment model was the top model for twilight and contained predictors based on land cover and cloud cover (Table 2; Figure 3). During twilight, we detected an interaction between land cover and cloud cover indicating a general decrease in deer movement selection through wetlands as cloud cover increased (85% CI –0.35, –0.13; SE = 0.07). Deer neither selected nor avoided paths though other land cover types based on cloud cover (Table 3). Based on our k-fold cross validation, both day and twilight models predicted selection better than would be expected at random. However, the top model for night (i.e., the global model; Table 2) failed to predict path selection better than would be expected at random despite containing informative parameters.

**Table 2.** Top models for predicting path selection for white-tailed deer (*Odocoileus virginianus*) during day, night, and twilight time periods in Central Florida, August – September 2019. Abbreviations are as follows: model, refers to the hypothesis evaluated; K, number of parameters; logLik, maximum log-likelihood; ΔAICc, difference of Akaike’s information criterion (AIC) between a model and the model with the smallest AIC; weight, model weight.

| Time period | Model | K | logLik | $\Delta AIC$ | Weight |
| --- | --- | --- | --- | --- | --- |
| Day | Forest Light Environment | 3 | -19465.8 | 0.0 | 0.74 |
|  | Thermal Refuge | 6 | -19464.1 | 2.6 | 0.20 |
|  | Dynamic Illumination | 6 | -19465.5 | 5.3 | 0.05 |
|  | Global | 17 | -19456.4 | 9.2 | 0.01 |
|  | Converging Light Environment | 8 | -19468.2 | 14.9 | 0.00 |
| Twilight | Converging Light Environment | 8 | -3001.6 | 0.0 | 0.88 |
|  | Forest Light Environment | 3 | -3008.8 | 4.5 | 0.10 |
|  | Thermal Refuge | 6 | -3007.8 | 8.5 | 0.01 |
|  | Dynamic Illumination | 6 | -3008.1 | 9.0 | 0.01 |
|  | Global | 17 | -2998.5 | 12.3 | 0.00 |
| Night | Global | 34 | -12584.2 | 0.0 | 1.00 |
|  | Converging Light Environment | 16 | -12618.4 | 32.0 | 0.00 |
|  | Forest Light Environment | 6 | -12628.9 | 33.0 | 0.00 |
|  | Thermal Refuge | 12 | -12625 | 37.1 | 0.00 |
|  | Dynamic Illumination | 12 | -12627.4 | 42.0 | 0.00 |

**Figure 2.**
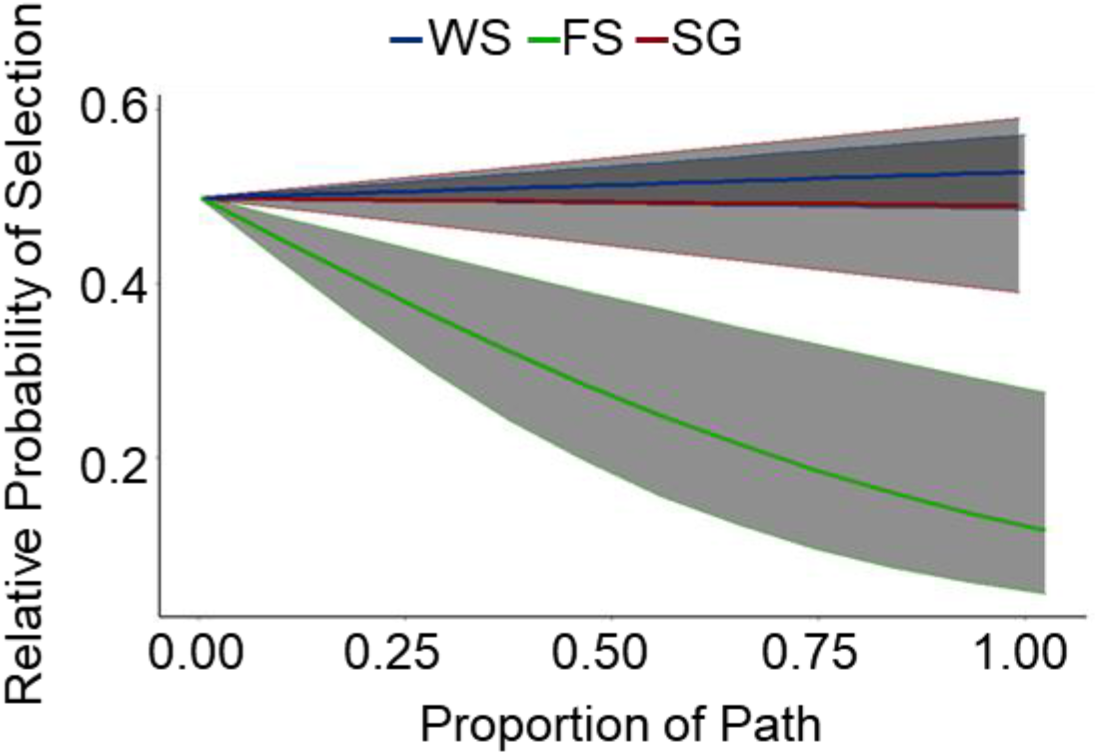
Relative probability of a white-tailed deer (*Odocoileus virginianus*) selecting a path during the day through woodland shade (WS), forest shade (FS), and small gap (SG) in Central Florida, August – September 2019. The shaded region represents confidence intervals where alpha = 0.5.

**Figure 3.**
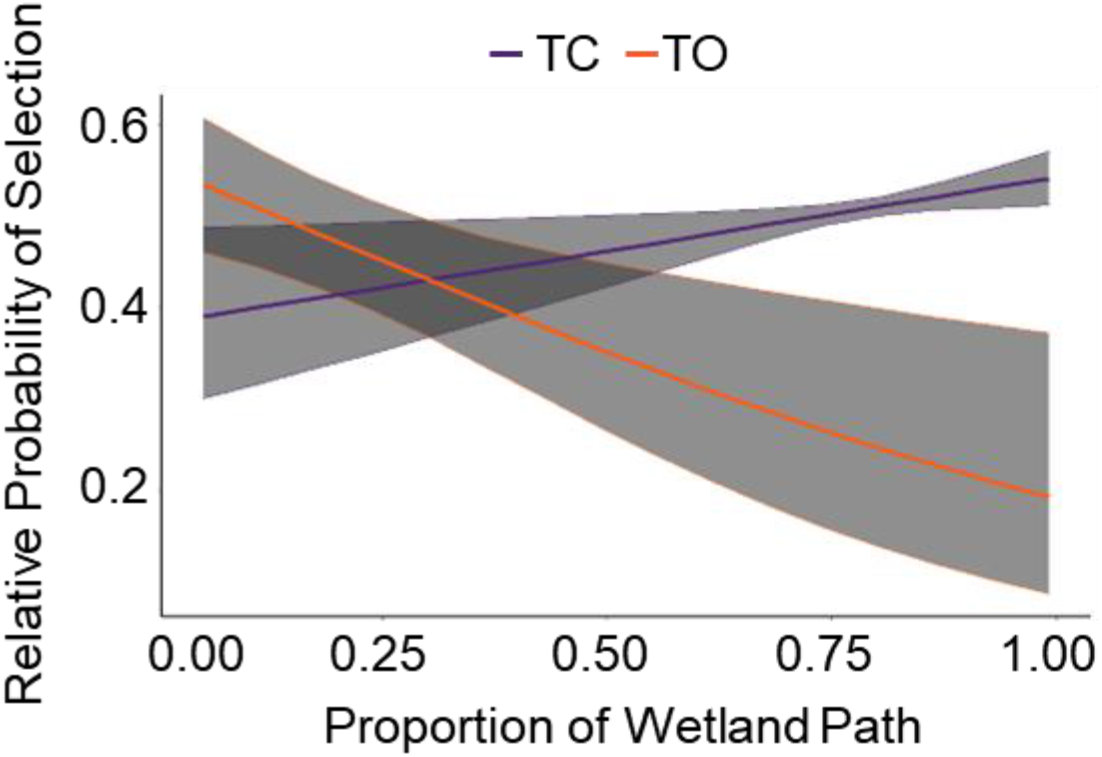
Relative probability of a white-tailed deer (*Odocoileus virginianus*) selecting a path through wetlands during twilight based on if cloud conditions were clear (TC) or overcast (TO) in Central Florida, August – September 2019. The shaded region represents confidence intervals where alpha = 0.5.

**Table 3.** Parameter estimates, standard error (SE), and 85% CIs for the top model predicting path selection for white-tailed deer (*Odocoileus virginianus*) during twilight time periods in Central Florida, August – September 2019.

| Predictor | Estimate | SE | Lower 85% CI | Upper 85% CI |
| --- | --- | --- | --- | --- |
| Wetland | 0.03 | 0.08 | -0.08 | 0.14 |
| Developed | 0.04 | 0.10 | -0.11 | 0.19 |
| Forested | 0.12 | 0.06 | 0.04 | 0.21 |
| Shrub and herbaceous | 0.00 | 0.06 | -0.08 | 0.09 |
| Wetland*cloud cover | -0.24 | 0.08 | -0.35 | -0.13 |
| Developed*cloud cover | -0.02 | 0.11 | -0.18 | 0.14 |
| Forested*cloud cover | -0.01 | 0.05 | -0.08 | 0.06 |
| Shrub and herbaceous*cloud cover | -0.03 | 0.06 | -0.12 | 0.05 |

## DISCUSSION

Thermal refuge, dynamic illumination, and land cover alone were not informative predictors of white-tailed deer movement decisions in Central Florida, rather light environments influenced these decisions during the day and at twilight. During the day, bucks selected to move through and avoid different forest light environments. For a signal or cue to be seen, it must first be detectable. Both spatial resolution and temporal resolution improve with increasing contrast and light levels (Donner, 2021; Peichl, 2005). Selection for woodland shade by white-tailed deer during movement might be related to the relatively high-light levels of this forest environment and short-wavelength enrichment compared to forest shade (Endler, 1993). Other ungulates rely on vision to distinguish high- and low-quality forage (Hirata and Kusatake, 2020; Hirata and Kusatake, 2021) and potentially locate foraging patches from long distances using visual cues (Bergman et al., 2005). The ability of white-tailed deer to discriminate details of their environment, including potential forage patches, would be improved within the high-light levels of woodland shade. Additionally, the short-wavelength enriched woodland shade environment in conjunction with the short-wavelength sensitivity of white-tailed deer might provide ideal conditions for intraspecific communication. Both caribou and elk (*Cervus elaphus*) pelage reflect short wavelengths at values higher than vegetation (Leblanc et al., 2016; Terletzky et al., 2012). If white-tailed deer pelage has similar spectral characteristics, conspecifics might be generally more conspicuous in woodland shade than forest shade. Furthermore, Caro et al. (2020) suggested both white-tailed deer and black-tailed deer (*O. hemionus*) use flash behaviors to confuse predators and avoid capture. Only conspicuous flash displays effectively reduce predation risk (Bae et al., 2019). Thus, increased contrast in woodland shade might make the flash behavior of white-tailed deer tail-flagging more effective as a distraction tactic for predators, as well as a threat signal for conspecifics.

Tradeoffs between signaling and unintended interspecific cueing of predators likely exist in woodland shade since predators of white-tailed deer are also sensitive to short wavelengths (Amann et al., 2014; Cronin and Bok, 2016; Douglas and Jeffery, 2014). Middle-wavelength enrichment in forest shade overlaps with the spectral sensitivity of white-tailed deer, however, spectral contrast between visual signals is likely reduced in forest shade relative to woodland shade. Reduced contrast in forest shade would decrease the effectiveness of flash behaviors for white-tailed deer, but interspecific visual cues to predators would also be limited in forest shade. White-tailed deer can exhibit both sexual and seasonal differences in resource selection and movement patterns (Beier and McCullough, 1990; Webb et al., 2010). While we observed an avoidance of forest shade during movement, deer might select to move through forest shade at different times of the year as ecological pressures change or select for this environment during other behaviors (e.g., bed-site selection) because of decreased conspicuousness in forest shade.

White-tailed deer, and other cervids, detect and respond behaviorally to perceived differences in predation risk (Cherry et al., 2015; Crawford et al., 2021; Gulsby et al., 2018; Price et al., 2014). At twilight, white-tailed deer in our study avoided wetlands during cloud cover. In the southeastern U.S., American alligators (*Alligator mississippiensis*) predate both juvenile and adult white-tailed deer (Shoop and Ruckdeschel, 1990) and more frequently attack prey during twilight and nocturnal hours (Nifong et al., 2014). The alligator population numbers in the thousands within the St. John’s River and its adjoining lakes (A. M. Brunell, Florida Fish and Wildlife Conservation Commission, personal communication) and threat of alligator predation is probable for white-tailed deer in wetlands. A spatiotemporal shift in the landscape of fear (Gaynor et al., 2019; Palmer et al., 2017) for white-tailed deer might occur due to the long-wavelength enrichment with cloudy weather during twilight. White-tailed deer are less sensitive to long wavelengths with a sharp decline in sensitivity at wavelengths > 600 nm (Cohen et al., 2014; Jacobs et al., 1994), while alligators have a long-wave sensitive cone with a peak sensitivity of 566 nm and minimal drop-off at wavelengths > 600 nm (Shoop and Ruckdeschel, 1990; Sillman et al., 1991). Thus, reduced visual perception by white-tailed deer might lead to the avoidance of wetlands during twilight under these specific light conditions and given risk of alligator predation.

In general, conflicting evidence exists regarding the influence of lunar illumination on white-tailed deer activity and resource use. Beier and McCullough (1990) found moonlight had little or no influence on the activity and resource use of white-tailed deer, while Newhouse (1973) found decreased use of open habitats on moonlit nights, and Brown et al. (2011) found increased use of open habitats on moonlit nights. We think our failure to successfully predict nocturnal path-selection of deer during movements based on light environment is unsurprising. The nocturnal forest landscape is primarily dominated by middle wavelengths (peak of 560 nm) with only small relative changes in light environment associated with lunar illumination, forest geometry, and cloud cover (Veilleux and Cummings, 2012). We suggest the visual specializations of white-tailed deer (Cohen et al., 2014; D’Angelo et al., 2008; Jacobs et al., 1994) might make seeking out these small relative changes in nocturnal light environments across the landscape unnecessary during movements. However, it is possible that a relationship might exist between lunar illumination and the nocturnal movement decisions of white-tailed deer at other times of the year. Lunar altitude and azimuth vary throughout the year due to the axial tilt of the earth and moon in their orbits, and the moon’s location in the sky has a greater influence on total lunar illumination than the phase of the moon (Todd et al., 2015; Veilleux and Cummings, 2012). Consequently, lunar illumination might play a more influential role in the nocturnal movement decisions of white-tailed deer during different seasons (e.g., winter vs summer).

Animals often possess visual specializations optimized for their ecology, and movement decisions should vary depending on these species-specific specializations. For white-tailed deer, habitat types of varying forest geometry might be important because the perceptual advantages and disadvantages of these environments are temporally variable and, in some cases, might be dependent upon activity (e.g., encamped vs movement) and predation risk (e.g., a long-wave sensitive predator). Our results highlight the need to consider light environments as a component of the multi-dimensional niche space used by white-tailed deer to meet their ecological needs. Additionally, when we think about vegetative requirements for this ungulate prey species, we likely should consider more than food, concealment, and thermoregulatory properties but also light environments. In conclusion, our findings suggest visual perception relative to light environments is likely an underappreciated driver of behaviors and decision-making by an ungulate prey species. Further investigation to understand the relative importance of light environments across various ecosystems and ungulate species would enhance our understanding of species-specific behaviors and their broader ecological relationships.

## ACKNOWLEDGEMENTS

We thank SITKA^®^ Gear for their support and desire to advance the knowledge base of wildlife visual systems. We thank the Florida Fish and Wildlife Conservation Commission for their help with permitting and obtaining funding. We thank J. Field, J. Smith, J. Taylor, C. R. Morea, R. M. Peters, and E. P. Garrison for their contributions for a safe and efficient project. Finally, we thank the landowners for their generosity and support of this study.

## COMPETING INTERESTS

Authors have no competing interests to declare for this work.

## FUNDING

This study was supported by SITKA^®^ Gear and the Fish & Wildlife Foundation of Florida, Inc. through the Wildlife Foundation of Florida Tag grant program (WFF 1819-17).

## DATA AVAILABILITY (if accepted)

Analyses reported in this article can be reproduced using the data provided by Newman et al. (2022).

**Newman, B. A., Dyal, J. R., Miller, K. V., Cherry, M. J., D’Angelo, G. J**. (2022). Data from: Influence of visual perception on movement decisions by an ungulate prey species. *Journal of Experimental Biology*. http://link to data.

## Notes

### Competing Interest Statement

The authors have declared no competing interest.

